# GenoPipe: identifying the genotype of origin within (epi)genomic datasets

**DOI:** 10.1101/2023.03.14.532660

**Authors:** Olivia Lang, Divyanshi Srivastava, B. Franklin Pugh, William KM Lai

## Abstract

Confidence in experimental results is critical for discovery. As the scale of data generation in genomics has grown exponentially, experimental error has likely kept pace despite the best efforts of many laboratories. Technical mistakes can and do occur at nearly every stage of a genomics assay (i.e., cell line contamination, reagent swapping, tube mislabelling, etc.) and are often difficult to identify post-execution. However, the DNA sequenced in genomic experiments contains certain markers (e.g., indels) encoded within and can often be ascertained forensically from experimental datasets. We developed the Genotype validation Pipeline (GenoPipe), a suite of heuristic tools that operate together directly on raw and aligned sequencing data from individual high-throughput sequencing experiments to characterize the underlying genome of the source material. We demonstrate how GenoPipe validates and rescues erroneously annotated experiments by identifying unique markers inherent to an organism’s genome (i.e., epitope insertions, gene deletions, and SNPs).

## INTRODUCTION

Biological discoveries in the genome sciences are heavily driven by high throughput sequencing (HTS) (1-4). As sample parallelization continues to grow, so too does the risk of technical errors in sampling (5-7). Technical error can be introduced at nearly every step of an experiment ranging from the mislabelling of the initial experimental sample tubes to errors in bioinformatic processing (8). While careful planning and attention to detail can mitigate many of these issues, it remains an omnipresent concern in evaluating and trusting data (9-13). Stringent quality control metrics are a practical and scalable solution to the important challenge of ensuring the dissemination of quality, reproducible genomic datasets (14-16).

Guidelines set by the NIH recommend regular authentication of cultured cell lines to address these problems (17). Cell line authentication is typically performed using multiple assays including microscope-based inspection of cell morphology, karyotyping, Single Nucleotide Polymorphism (SNP) Profiling, and Short Tandem Repeat (STR) Profiling (9,18-21). STR profiling detects allelic differences within microsatellite repeat regions of cell lines and is based on the principal that each distinct cell lines possesses a unique and reproducible microsatellite variant signature. The American Type Culture Collection has set STR profiling as its recommended standard due to its high specificity and broad accessibility for most labs (12,22). Another suggested approach includes using DNA-barcoding technology coupled to whole-genome sequencing for authenticating cell lines (23). This demonstrated that genomic approaches can be used to validate a cell line’s identity, although regular whole-genome sequencing can be cost-prohibitive for many research projects.

While these approaches are excellent for validating stock cultures, for researchers performing downstream genomic experiments on those cell lines, they represent an additional assay to perform and therefore an additional avenue for sample metadata mislabelling. Additionally, recent advances in genome editing technologies (e.g., CRISPR, ZFNs, TALEN) have dramatically expanded the possibilities for engineering genomes directly to include specifically localized insertions and deletions (24-27). Genes of cell lines are often modified to contain an artificial epitope fused to the target protein for purposes including facilitating the purification of the gene product, fluorescently tagging the protein, or conditionally depleting the gene product (28-31). The application of this technology has been widespread in many areas of biological and genomic research (32). There are now many cell lines and strains that have been specifically engineered to contain epitope-tagged genes for the purposes of facilitating purification of the protein or simply to mutate the gene itself (28,33-36). Critically, these novel cell lines theoretically possess the same microsatellite profile as stock strains even though they are now genotypically distinct, making STR validation of limited value for genetically modified cell lines.

Validating cell line identity within each DNA sequencing dataset directly ties cell identity to each experimental dataset. This differs from a separate cell line authentication assay (i.e., STR profiling) which cannot rule out any downstream sample mix-up or contamination during any of the biochemical or bioinformatic processes taken to generate the aligned reads. Previous bioinformatic approaches have used Bayesian approaches to identify strain backgrounds but were limited to whole-genome sequencing (WGS) assays (37). Another approach (CeL-ID) combined the Genome Analysis Toolkit (GATK), VarScan, and the variant database COSMIC to characterize a cell line background in RNA-seq data (38-41). While this tool provides detailed and accurate cell line authentication, it is specific to RNA-seq assays which precludes its use on a wide variety of genomic assays.

We developed the <u>Geno</u>type validation <u>Pipe</u>line (GenoPipe) to calculate the unique molecular signature inherent within genomic data from a wide range of biochemical assay types in order to confirm sample metadata and potentially rescue improperly labelled samples. For variant-based backgrounds, reads that cover sites that occur within a known variant profile can be checked for signal supporting certain cell identities. These signals in the dataset provide an avenue for GenoPipe to perform quality controls that confirm a sample’s expected genetic background. Using GenoPipe, we have tested multiple large-scale genomic datasets (i.e., human ENCODE, yeast YKOC deletion) and have identified apparent errors in advertised sample identity. GenoPipe provides a mechanism to recover these samples and restore trust in genomic analysis using these sampels.

## MATERIAL AND METHODS

### Software Dependencies

Each module of GenoPipe is a distinct heuristic script that runs a series of custom Python and Perl scripts and commands of third-party software. These include Bowtie2, BWA-MEM (v0.7.14), bedtools (v2.26), samtools (v1.7), and popular python libraries like numpy, scipy, and pysam (42-47). For the DeletionID and StrainID modules that use aligned input BAM data, it is plug-and-play compatible with data using alternative aligners (e.g., BWA-MEM and bowtie2). The software can be download from GitHub (https://github.com/CEGRcode/GenoPipe) and includes extensive documentation for setup and usage.

### Reference Genomes and Annotations

All yeast data was aligned to the sacCer3 genome and gene annotations were downloaded from the Saccharomyces Genome Database (SGD) (https://www.yeastgenome.org/). The hg19 genome was downloaded from UCSC Genome Browser (http://hgdownload.cse.ucsc.edu/goldenPath/hg19/bigZips/hg19.2bit) and gene annotations were downloaded from RefSeq (ftp://ftp.ncbi.nlm.nih.gov/refseq/H_sapiens/annotation/GRCh37_latest/refseq_identifiers/GRCh37_latest_genomic.gff.gz). Scripts for downloading the genomes and annotations and for building the GenoPipe reference files for each module are included in the Github repository within the respective ‘utility_scripts’ directories.

### EpitopeID algorithm

The EpitopeID module is executed with scripts that run Bowtie2, bedtools, samtools, and custom Perl5 and Python scripts. The reference files required by EpitopeID are a genomic FASTA sequence, gene annotations GFF file, and tag or epitope FASTA sequences. EpitopeID provides pre-built reference files for yeast (sacCer3) or human (hg38/hg19), to update the tag database with new tag sequences, and to convert the set of GFF gene annotations into a EpitopeID reference GFF file of labelled genomic bins. The default genome binning references create a bin for the length of each gene coding region, an upstream “Promoter” bin 250bp in size, a downstream “C-term” bin 250bp in size, and the rest of the intergenic regions broken up into bins of 250bp.

The main ‘identify-Epitope.sh’ script begins by aligning all FASTQ reads to the epitope sequences in the tag database. For both single-end and paired-end data, the module will report the number of reads that map to each tag in the tag database.

If the data is paired-end, the module will then align the mate-pair of these reads to the genome. Reads that map to unwanted or artefactual regions of the genome are removed (i.e., blacklist filter is applied to the set of reads). The 5’ ends of the mapped reads are tallied for each genomic bin (*i*) to get a read count (*k*_*i*_). Using the read count (*k*_*i*_) and bin size in bp (*b*_*i*_), the genome size in bp (*G*), and the user-provided fold-minimum enrichment (*M* defaults to 2), the tool calculates the probability (*λ*_*i*_) of a single read falling in each bin *i*.

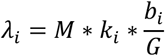

Using these probabilities (*λ*_*i*_), and the read counts (*k*_*i*_), the tool calculates a p-value score (*p*_*i*_) for each genomic bin *i* based on a poisson distribution.

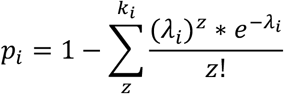

This p-value score represents the chance of having the observed read count or greater for the genomic bin *i*. The module will report all bins with p-value scores over the user-provided threshold sorted from most significant to least.

### DeletionID algorithm

The DeletionID module is executed with scripts that run BWA-MEM, bedtools, and custom Perl and Python scripts. The genomic interval reference files required by DeletionID along with the utility scripts used to generate them are available at: ‘GenoPipe/DeletionID/utility_scripts’.

The ‘sacCer3_Del’ yeast genomic intervals are based on the SGD ORF coordinates (http://sgd-archive.yeastgenome.org/curation/chromosomal_feature/saccharomyces_cerevisiae.gff.gz) for all Verified, Uncharacterized, Dubious, and Blocked Reading Frame ORFs. Each genomic interval was saved as a BED coordinate file and a “mappability” score for each interval is calculated. These mappability scores (*M*_*ij*_) are determined by measuring the ratio of for each coordinate interval *i* and read length *j*. The mappability score is calculated by first tiling the given coordinate interval *i* by various read lengths *j* (100, 36, 40, and 50 base pairs) every 25bp. The sequences of each tile are extracted and then mapped to the whole genome. These tiles are filtered to keep the uniquely mapping reads. The mappability score (*M*_*ij*_) of coordinate interval *i* and read length *j* is the number of uniquely mapped reads (*u*_*ij*_) scaled by the step size (default: 25 base pairs) and normalized to the interval size (*l*_*i*_) to make it a mappability per bp score.

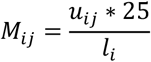

Using these mappability scores as a reference, a score (*S*_*i*_) is calculated for each coordinate interval i using the interval size (*l*_*i*_), the mappability of the coordinate interval (*M*_*ij*_), and

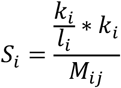

The set of scores is filtered to exclude the coordinate intervals that don’t meet a user-defined mappability threshold (defaults to filter intervals with *M*_*i*_ < 0.25). The final score (*F*_*i*_) takes the filtered scores (*S*_*i*_) and scales them all down by the median of the set of filtered scores and puts them in log space for each coordinate interval *i*.

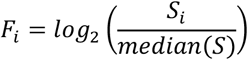

Each coordinate interval that was not filtered out is printed in the report file with its final score.

### StrainID algorithm

The StrainID module is executed with scripts for a custom python script that performs the retrieval of BAM reads and FASTA genomic sequence information, tallies up the number of alternate and reference alleles for each VCF variant profile and the background signal score to calculate the final log score. Recommended input VCF variant profile databases for StrainID are available on GitHub for some popular strains and cell lines in yeast and human based on the sacCer3 and hg19 genome builds.

The ‘sacCer3_VCF/full_VCF’ files were generated by Song et al and downloaded via SGD (48). This set includes 11 commonly used strains in yeast research and industry. The recommended and default VCF database for sacCer3 data provided by GenoPipe was constructed by taking the subset of each variant profile to include only the variants that are unique to each profile among these 11 strains.

The ‘hg19_VCF’ files contain the variants for 8 cell lines: A549, HCT116, HELA, HepG2, K562, LnCap, MCF7, and SKnSH. These variants are based on the hg19 reference genome build which is the human genome build used for processing human data in StrainID. They also include only exomic variants, or variants that fall within the gene annotations for the reference genome.

A variant score (*S*_*i*_) is independently calculated for each variant profile (*V*_*i*_) within the VCF database (e.g., ‘sacCer3_VCF’ or ‘hg19_VCF’) according to the following method.

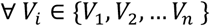

For each substitution variant *v* in a given variant profile *V*_*i*_ that consists of a set of variants (i.e., VCF), the reads that align to the variant site are identified and tallied into two counts: the number of reads that show the alternate allele (*a*_*v*_) and the number of reads that show the reference allele (*r*_*v*_) for some variant *v*. A background model is built by sampling either 1 million or 5% of mapped reads, whichever is greater, and tallying up the number of substitution mismatches found in each read (*b*_*alt*_) and tallying the number of nucleotides that match the reference (*b*_*ref*_). If the number of mapped reads is less than 1 million, then the whole dataset with all mapped reads is used for the background model. The following equation is used to calculate a final StrainID score for each variant profile.

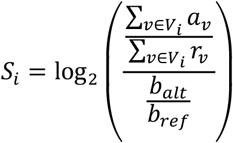

The largest StrainID score value of the output report indicates the predicted strain background.

### Synthetic Depth simulations

Each module was tested against simulated data generated from synthetic genomes that were created using scripts available on Github to simulate random synthetic epitope insertions, gene deletions, and SNP-replacement of reference alleles for alternate alleles.

The depth-test simulations were generated by sampling coordinates for the paired end reads that are each 40bp long and 100-350 bp apart (uniform sampling of distance between read pairs). The genomic FASTA from the *in silico*-modified genomes were pulled at these coordinates to generate a FASTQ file with fixed Phred scores across all read positions for all reads. EpitopeID runs on these FASTQ files directly. BAM files for running StrainID and DeletionID tests were created by aligning to the sacCer3 and hg19 reference genomes using BWA-MEM (42), then sorting and indexing the BAM using samtools (47).

### Synthetic Epitope Contamination Simulations

The contamination simulations for EpitopeID were generated by sub-setting the simulated FASTQ files from the depth simulations to 9 sizes: 90 thousand reads decrementing down to 10 thousand reads by 10 thousand read increments. The RAP1-R500 subsets were concatenated with the Reb1-R500 subsets to create 9 FASTQ files of 1 million reads each. The human simulations were similarly mixed for the CTCF-R500 background and the POLR2H-R500 background. These were all run through EpitopeID and genomic loci with significant hits were tallied to determine which contaminated datasets identified a REB1 locus and which identified a RAP1 locus.

Scripts for generating and running simulated data through each of the GenoPipe modules can be found in the GenoPipe GitHub repository under the ‘paper/SyntheticEpitope’ directory.

### ENCODE data selection and testing

Data accessions were identified using the ENCODE RESTful API to identify samples appropriate for testing EpitopeID and for testing StrainID (49). Separate python scripts were written for EpitopeID and StrainID testing to parse out the samples fitting each set of requirements. Sample accessions were obtained on February 27, 2021 for the epitope-tagged samples and May 12, 2021 for the human cell line samples.

For the EpitopeID datasets, all Genetic Modification accessions (ENCGM) were searched for those with the purpose of “tagging” and in the “insertion” category. Biosamples (ENCBS) associated with these Genetic Modification accessions were filtered for “Homo sapiens” as the organism. The biosample accessions were used to identify libraries (ENCLB) and then the raw FASTQ files (ENCFF) that were then downloaded and run through the EptiopeID module using the hg19 genome build and annotations. The “FASTA_Tag” database included the eGFP-containing LAP-tag, the 3xFLAG sequence, and the random 500bp sequence used for the epitope insertion simulations. The reported successes and failures for EpitopeID only included datasets with a “released” status and only the eGFP insertion modification.

For the StrainID datasets, all biosample accessions (ENCBS) with the type “cell line” were searched for those whose background matched one of the cell lines that StrainID uses in its default reference VCF database (i.e., A549, HCT116, HeLa, HeLa-S3, HepG2, K562, LNCAP, MCF-7, and SK-N-SH). Then all File accessions (ENCFF) were filtered to include the “bam” type files aligned to the hg19 genome build and whose biosample accession matched the above obtained ENCBS codes. These files were downloaded and StrainID was run against them using the default ‘hg19_VCF’ database.

### Identifying genomic HIV insertions

The hg19_EpiID database was updated with the HIV genomic sequence (GenBank accession AF324493.2, HIV-1_vector_pNL4-3) in the tagDB directory. Raw FASTQ files of ChIP-seq data performed in HIV-infected T-cells were downloaded from SRA and can be accessed through the GEO accession GSE84199 (50). EpitopeID was run on these datasets using this custom hg19_EpiID database.

### YKOC analysis

The 9010 sequencing datasets of whole genome sequencing (WGS) of the YKOC samples were downloaded from the EBI project accession PRJEB27160 (51). The list of ERR run accessions used are listed with the results in **Supplementary Table 2**. The raw FASTQ files were aligned to the sacCer3 genome using BWA-MEM. The resulting BAM files were sorted and indexed and run through DeletionID using the same ‘sacCer3_Del’ database used for the yeast simulations. DeletionID was run with default parameters, filtering genes for a mappability greater than 25% and filtering the on a log2 output threshold of less than -2 for the final scores.

### Yeast ChIP-seq analysis

The 14 ChIP-seq sequencing datasets (four CENPK background and 10 BY4742 background) were downloaded using their respective SRA accessions (52,53). The Illumina datasets were aligned with BWA-MEM using default parameters on the sacCer3 genome build while the ABI SOLiD datasets were aligned with Bowtie (v1) using a color space reference index built from the sacCer3 genome build. The sorted and indexed BAM files were then run through StrainID using the provided ‘sacCer3_VCF’ database and the results were parsed into a table for visualization.

## RESULTS

### GenoPipe design and implementation

GenoPipe is composed of three analysis modules, each designed to parse high-throughput sequencing datasets to identify specific classes of alterations relative to a reference genome. The <u>Epitope</u>-tag <u>ID</u>entification (EpitopeID) module searches for the presence of known DNA sequences (e.g., synthetic protein epitopes) in a FASTQ file and then leverages paired-end sequencing to identify the site of insertion in the host genome based on the alignment of the mate-pair (**Figure 1A**). The <u>Deletion ID</u>entification (DeletionID) module models the background of a genomic experiment to identify significantly depleted regions of the genome to predict genomic deletions (**Figure 1B**). It then looks up what annotated features overlap the deletion. The <u>Strain ID</u>entification (StrainID) module uses existing SNP or variant calls databases of common cell lines (K562, MCF7, HepG2, etc.) to match a cell’s genetic identity inherent to each experimental NGS dataset to the set of variant calls that are unique to each cell line (**Figure 1C**).

**Figure 1.**
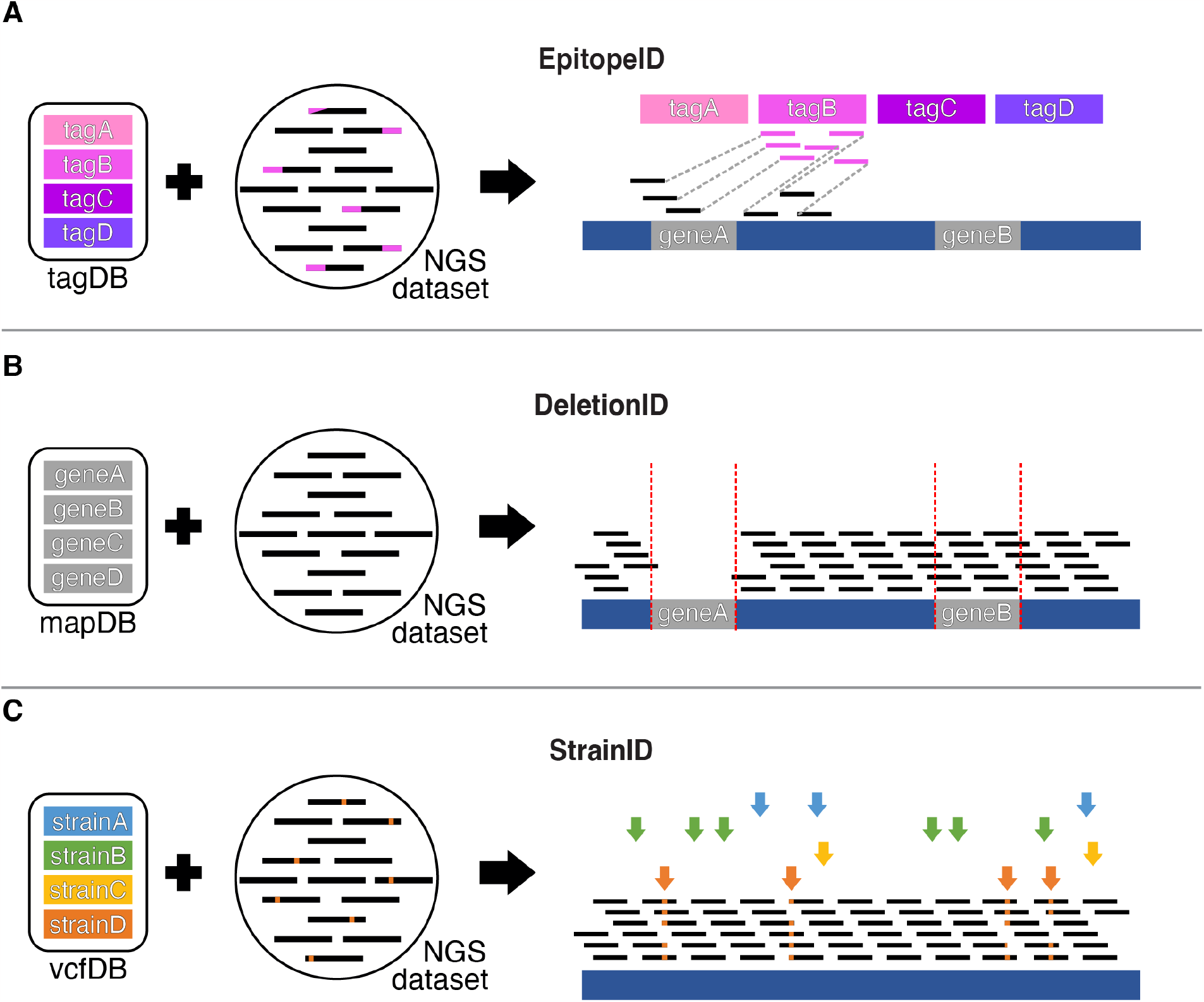
Schematic of the GenoPipe’s three modules. (**A**) The EpitopeID module identifies known epitope tag insertions by aligning raw reads (NGS dataset) to a set of tag sequences (tagDB). It then localizes the tag by mapping the mate-pair of tagDB-aligned reads to a reference genome and looking for enrichment of annotated features (e.g., geneA) around the insertion site. (**B**) DeletionID looks for annotated gene intervals (mapDB) with depleted genomic alignments to identify deletions (e.g., geneA) in the strain background of a NGS dataset. (**C**) StrainID identifies a variant-based strain background (vcfDB) that best matches a given NGS dataset (colored arrows) by assigning a score for each Variant Call File (VCF) provided by a user.

### EpitopeID Module: Genomic sequence identification and localization

The EpitopeID module identifies the presence and approximate location of specific DNA sequences within the genome. The algorithm functions by first aligning the raw sequencing data (i.e., FASTQ) against a curated DNA sequence database (tagDB) of common protein epitopes provided by the tool that is easily customized by the user to include other “tag” sequences (**Figure 1A**). EpitopeID reports the alignment statistics to all DNA sequences in tagDB. When the sequencing data exists as paired-end, EpitopeID will also align the mate-pair of reads that previously aligned to the tagDB to the reference genome. This provides the approximate genomic location of the epitope sequence and its probable fusion partner.

The EpitopeID algorithm was initially developed on simulated sequencing data from 18 *in silico*-modified genomes, each containing an artificial epitope insertion inserted at a different target locus in a genome. The three epitopes tested were one random sequence 500 bp long (R500), one random sequence 100 bp long (R100), and one random sequence 50 bp long (R50). These were each inserted into the N-terminus or C-terminus of three genes (*RAP1, REB1*, and *SUA7*) in the 12 megabase (MB) budding yeast genome (sacCer3) and of three genes (CTCF, POLR2H, and YY1) in the 3 gigabase (GB) human genome (hg19). Paired-end FASTQ sequences were randomly sampled assuming a uniform distribution from these *in silico*-modified genomes at varying levels of theoretical sequencing depth. The simulated sequencing files were then used as input for the EpitopeID system. In our simulations on the yeast genome using only 100 thousand (K) unique paired-end tags, the correctly tagged gene was identified 97.6%, 96.5%, and 97.9% of the time for *RAP1, REB1*, and *SUA7* (**Figure 2A; Supplementary Table 1**). For human data, a sequencing depth of 20 million (M) unique paired-end tags identified the R500 epitope at the correct locus 93.4%, 95.4%, and 94.7% of the time for the CTCF, POLR2H, and YY1 loci, respectively (**Figure 2B; Supplementary Table 1**). These thresholds (100K in yeast and 20M in human) represent the recommended minimum sequencing depth required for EpitopeID to successfully identify the DNA insertion 500bp long in most cases for each respective organism. This sequencing depth matches the ENCODE recommended sequencing depth for mammalian ChIP-seq experiments as well as the minimum recommended sequencing depth for ChIP-based experiments in yeast (14,54). EpitopeID runtime is exceptionally efficient and sensitive as it relies on the highly optimized Bowtie2 alignment algorithm (43). EpitopeID runs to completion in ∼3 seconds for 100K simulated yeast reads and ∼4 minutes for 20M simulated human reads (**Figure 2C-D; Supplementary Table 1**).

**Figure 2.**
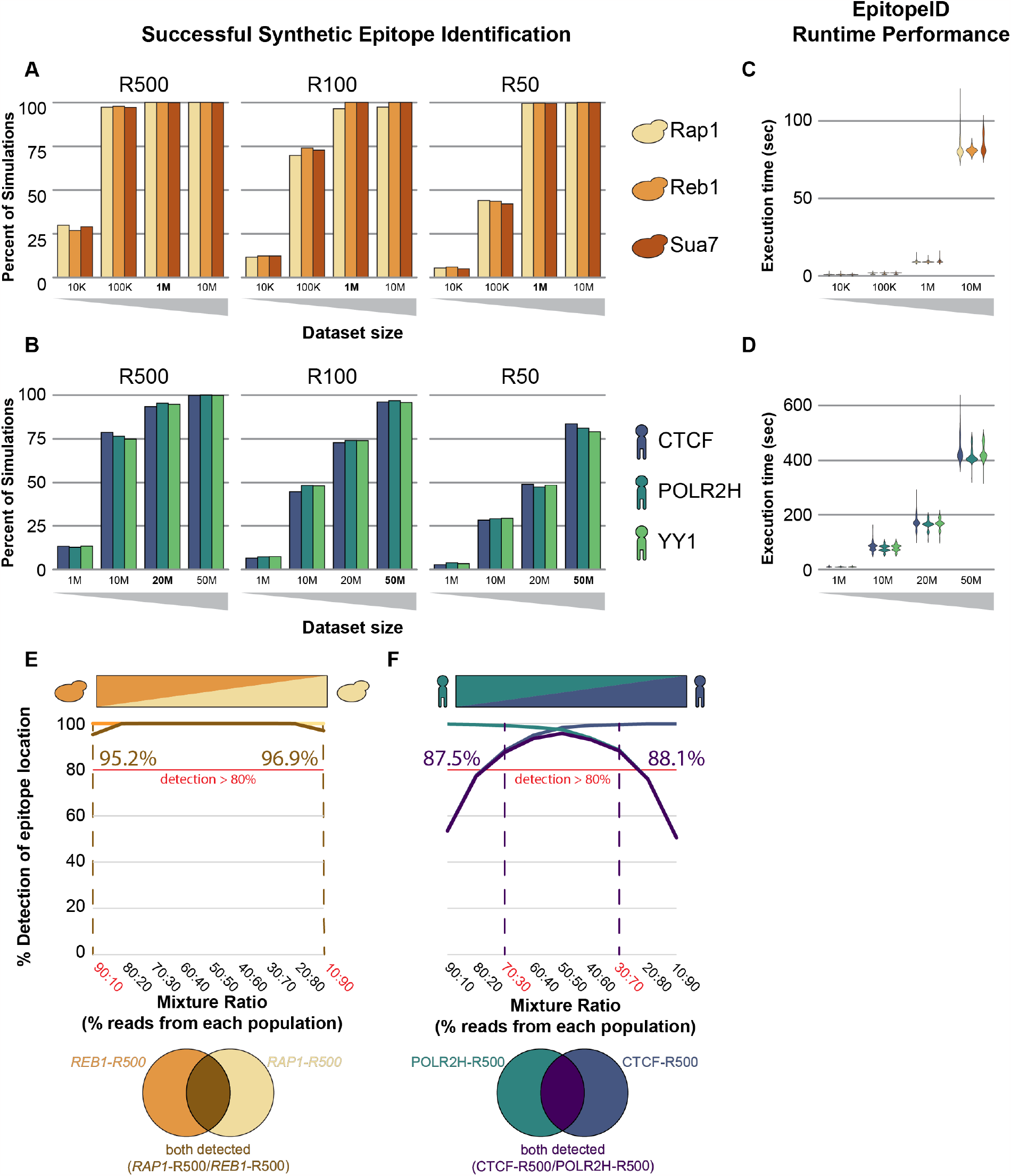
Evaluation of the EpitopeID module for sensitivity and performance. (**A**) Proportion of 1,000 simulations where EpitopeID successfully identified each of the variable length synthetic epitopes: R500, R100, and R50. Different colors represent different gene targets these epitopes were localized to (yellow:*Rap1*, orange:*Reb1*, and red: *Sua7*) in the yeast genome at multiple simulated sequencing depths. (**B**), Proportion of 1,000 simulations where EpitopeID successfully identified the synthetic epitope (R500, R100, and R50) and localized it to the correct target (dark blue:*CTCF*, blue:*POLR2H*, and green:*YY1*) in the human genome at multiple simulated sequencing depths. (**C**,**D**), Average runtime performance of EpitopeID for each of the R500 simulation sets in respective panels (**A)** and (**B)** across different sequencing depth in yeast (**C**) and human (**D**). (**E**), Proportion of epitopes detected when reads from 100K simulated sequence reads from *REB1*-R500 and *RAP1*-R500 were subsampled and mixed in titrating ratios(x-axis), and subsequently run through EpitopeID. The orange line represents the proportion when EpitopeID detected both *REB1* and *RAP1*. (**F)**, Proportion of epitopes detected when reads from 50M simulated sequence reads on CTCF-R500 and POLR2H-R500 were subsampled and mixed in titrating ratios(x-axis), were run through EpitopeID. The teal line represents the proportion when EpitopeID detected both CTCF and POLR2H.

We also examined the ability of EpitopeID to detect cellular cross-contamination by mixing reads simulated from two similar genotypes (same epitope tag inserted at two different loci) and counting how often each genotype was detected at different levels of contamination. In the yeast simulations, the *REB1*-R500 and *RAP1*-R500 in silico-modified genetic backgrounds were mixed at 9 levels of contamination (10, 20, 30, …, and 90% contamination) for a total of 1M reads (**Figure 2E**). For human, reads from the CTCF-R500 and POLR2H-R500 in silico-modified genetic backgrounds were mixed at the same 9 titrating percentages of contamination but with a total of 50M reads (**Figure 2F**). As expected, the ability of EpitopeID to detect a contaminating genome increased with increasing percentages of contamination. Using a threshold of detecting both backgrounds 80% of the time, we found that for yeast samples with 1M paired reads, EpitopeID can reliably detect both backgrounds in as little as 10% contamination while for human samples with 50M paired reads, both backgrounds can be reliably detected with a minimum of a little less than 30% contamination (**Figure 2E-F**; **Supplementary Table 2**).

### Identification of insertions in published data

We originally applied the EpitopeID heuristic to our comprehensive survey of DNA-binding proteins in the *S. cerevisiae* genome (54). Using EpitopeID we have guaranteed the identity of over 1,200 unique ChIP-exo datasets. We next moved to an examination of large collections of datasets available in the public domain.

We applied our EpitopeID system to 1,150 ChIP-seq and DNase-seq samples of paired and single end data generated by ENCODE (32). These samples contained a diverse set of eGFP and 3xFLAG-tagged proteins generated from various genome modification technologies including but not limited to: CRISPR and site-specific recombination across several cell lines, primarily K562, HepG2, and HEK293. We successfully confirmed the presence of at least one read mapping to the eGFP epitope in 1,144 of 1,150 datasets (99.5%) and at least one read mapping to the 3xFLAG epitope in 982 of 984 datasets (99.8%), regardless of whether its location matched the annotated location. As our simulations were based on paired-end data, we re-extrapolated the threshold for epitope detection in single-end sequencing to be 40M reads sequenced to achieve consistent coverage. All of the four single-end eGFP samples and both 3xFLAG samples that did not map to their respective epitope sequence failed to meet the required sequencing depth (20M paired-end, 40M single-end), and so remain undetermined. The remaining two paired-end samples sequencing depths were well beyond the simulation-based recommended minimum with somewhere between 20M and 50M reads. Based on the simulations we performed, there is between an 8.5% and 0.2% chance that the sequencing missed reads that would have mapped to the eGFP epitope. These samples should be investigated further for the possibility of a mislabelling or contamination problem.

For the 887 of the 1,150 datasets that contained both eGFP and paired-end reads as annotated by ENCODE, EpitopeID attempted to localize the epitope. Of these, 745/887 (84.2%) localized the tag to the expected gene target (**Table 1**). Two samples failed to localize the epitope because no reads mapped to eGFP and as a result, there were no mate pairs to map for localization information. Deeper investigation into some of these samples revealed suspicious metadata patterns that indicate likely sample mislabelling (**Figure 3**).

**Table 1.**
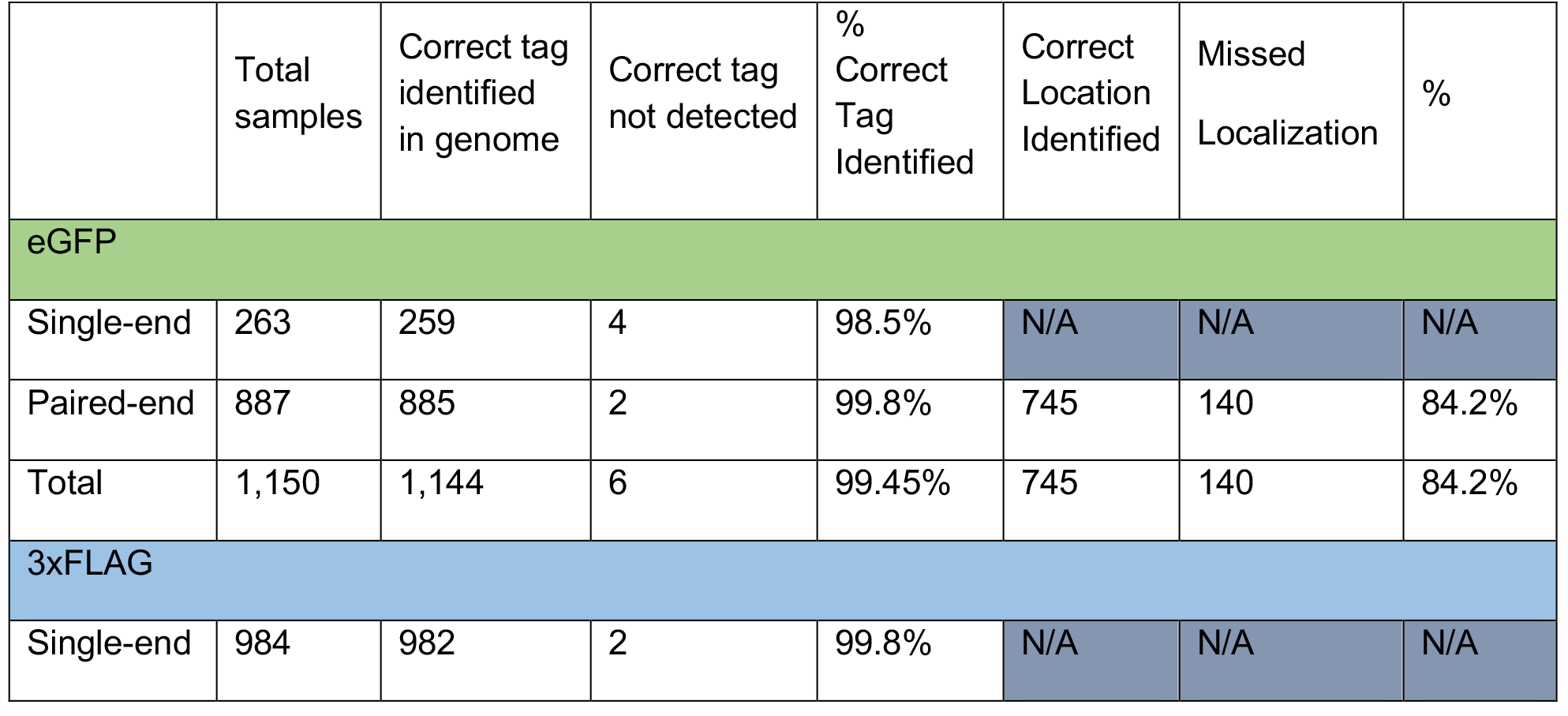
EpitopeID performance in identification and localization of epitopes in ENCODE data. The ENCODE datasets include both single-end and paired-end data with an ENCODE audit status of “released” from genetic backgrounds using the eGFP epitope and single-end data from genetic backgrounds using the 3xFLAG epitope. This table shows the number of samples in each category, the number of samples in which EpitopeID did or did not identify the correct eGFP or 3xFLAG epitopes respectively, the percentage of this correct identification, and for the paired end samples, whether or not EpitopeID correctly localized the epitope to the expected locus.

|  | Total samples | Correct tag identified in genome | Correct tag not detected | % Correct Tag Identified | Correct Location Identified | Missed Localization | % |
| --- | --- | --- | --- | --- | --- | --- | --- |
| eGFP |  |  |  |  |  |  |  |
| Single-end | 263 | 259 | 4 | 98.5% | N/A | N/A | N/A |
| Paired-end | 887 | 885 | 2 | 99.8% | 745 | 140 | 84.2% |
| Total | 1,150 | 1,144 | 6 | 99.45% | 745 | 140 | 84.2% |
| 3xFLAG |  |  |  |  |  |  |  |
| Single-end | 984 | 982 | 2 | 99.8% | N/A | N/A | N/A |

**Figure 3.**
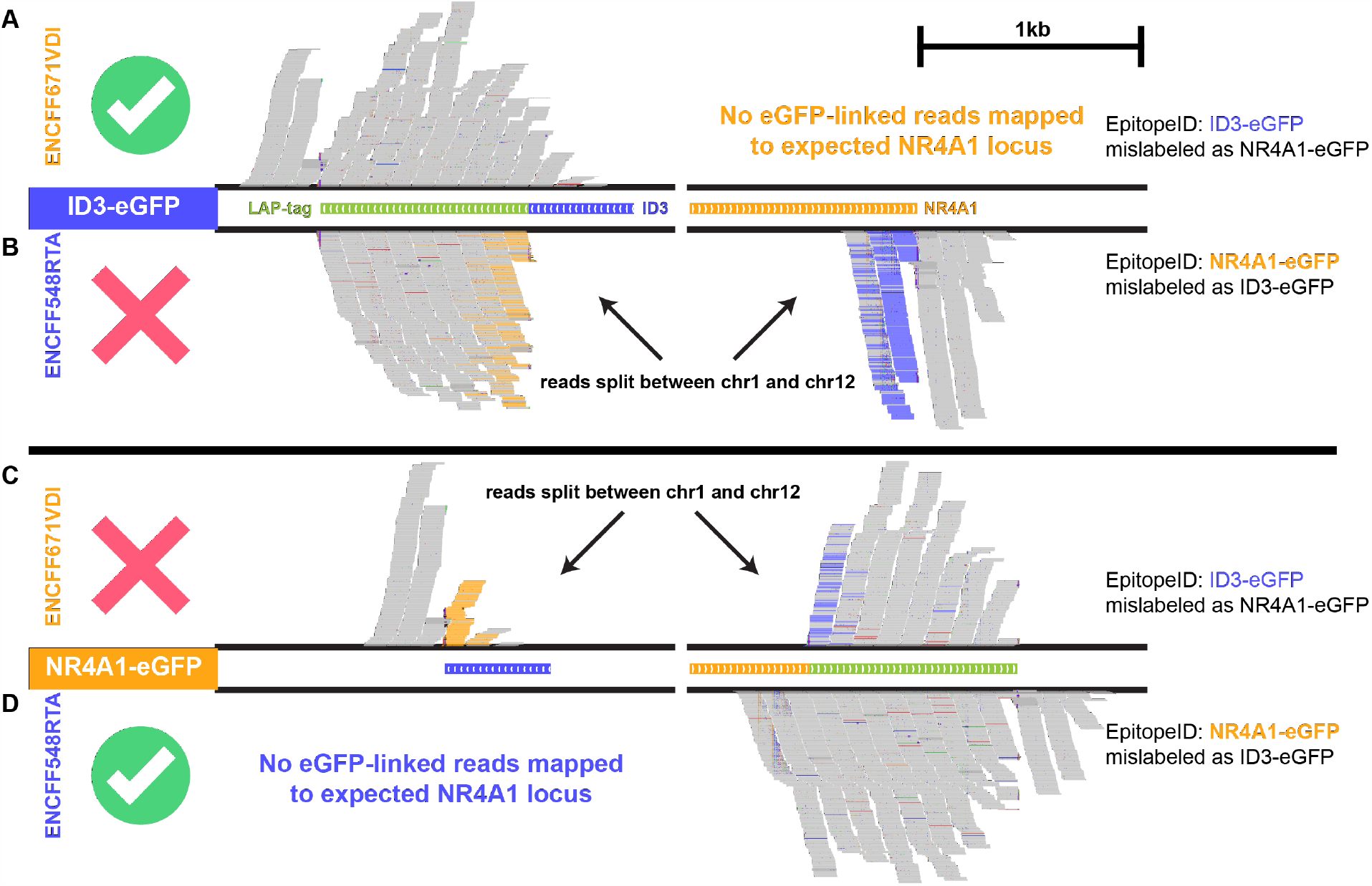
Alignment of two mislabelled ENCODE datasets whose genotype information is likely swapped. Genome browser shot of a subset of read pairs from two datasets where at least read maps to the LAP-tag (eGFP). Alignments were made to the ID3-eGFP genome for (**A**) an NR4A1-eGFP ENCODE labelled dataset and (**B**) an ID3-eGFP ENCODE labelled dataset, and aligned to the NR4A1-eGFP genome for the same (**C**) NR4A1-eGFP ENCODE labelled dataset and (**D**) ID3-eGFP ENCODE labelled dataset.

For example, six datasets that demonstrate metadata patterns and EpitopeID results consistent with a possible sample mix-up between an ID3-eGFP strain and an NR4A1 strain. Three of the six datasets are labelled with the ID3-eGFP genotype (ENCFF548RTA, ENCFF622ATI, ENCFF542CMJ) by ENCODE while the other three are labelled with the NR4A1-eGFP genotype (ENCFF671VDI, ENCFF236YCR, ENCFF760LCQ) by ENCODE. EpitopeID reported a strong NR4A1-eGFP genotype for all three ID3-eGFP labelled datasets and a strong ID3-eGFP genotype for all three NR4A1-eGFP datasets. To visually verify this in the genome browser, reads that align to the eGFP epitope and their mate pairs were each mapped to two modified reference genomes with the ID3-eGFP and NR4A1-eGFP genotypes. The reads that were labelled as coming from an NR4A1-eGFP strain showed continuous and unbroken mapping to the eGFP tag and flanking regions when it was inserted at the ID3 locus (**Figure 3A**) while the reads from a ID3-eGFP labelled sample showed discontinuous mapping between the eGFP sequence and the NR4A1 locus on a separate chromosome (**Figure 3B**). Conversely, when eGFP sequence is inserted at the NR4A1 locus and the reads are re-aligned to this new genome, the NR4A1-eGFP labelled sample showed discontinuous mapping between the eGFP sequence at the NR4A1 locus and the ID3 locus (**Figure 3C**) while the ID3-eGFP labelled sample showed full coverage of the eGFP locus and the flanking regions at the NR4A1 locus (**Figure 3D**). Other similarities such as submission timestamps from ENCODE suggest that this is the result of a simple mislabelling error during processing.

Outside of the handful of dataset clusters with these putative metadata exchanges, EpitopeID flagged recurring regions of off target eGFP localization across a number of unrelated datasets, especially localization to the HOXA13 and HOXB13 gene regions. It is difficult to determine at what stage this potential contamination is occurring since data collection is downstream of a long series of experimental steps, any of which could be causing the possible contamination. These “Hox contamination” datasets include a mix of both CRISPR-based and site-specific recombination tagging methods which suggests a tagging method-independent cause. One interesting observation in all of these datasets is that they were all generated using the Illumina HiSeq 4000 platform. Recent work has suggested that Illumina sequencers using pattern flow cell technology such as the HiSeq 4000 may incorrectly multiplex samples (55). This may explain the recurrent contamination signals that EpitopeID is detecting. A complete summary of each ENCODE eGFP ChIP-seq sample and the results of their analysis using the EpitopeID system is available in **Supplementary Table 3**.

In addition to performing quality control checks of data generated from samples with engineered genetic backgrounds, we also investigated EpitopeID’s ability to localize other sequence-based insertions. We applied EpitopeID to paired-end ChIP-seq data generated in HIV-infected T-cells (50). EpitopeID identified the presence of the HIV genome and successfully localized it to multiple integration sites in the genome (**Supplementary Table 4**). We found most of the integration sites EpitopeID identified were proximal to highly-transcribed protein encoding genes, which is consistent with previous work demonstrating HIV integration sites to be mainly within active transcriptional units (56). This result also indicates that EpitopeID can also serve for *de novo* discovery of viral insertions as well as identification and localization of genomic translocations.

### DeletionID Module: Identification of depleted genomic intervals

The DeletionID module identifies depletions of aligned reads in chromatin-based HTS data. It checks each user provided genomic interval and reports the intervals that have a depletion above a user-defined threshold. The algorithm functions by identifying aligned reads (i.e., BAM) within a set of genomic intervals and calculates a coverage score for each interval that factors in the read counts, mappability of the sequence in each interval (mapDB), and the length of the genomic interval (**Figure 1B**).

The DeletionID algorithm was developed by simulating HTS data from two in silico-modified genomes, each with a genomic interval removed. The *RAP1* and *REB1* ORFs were removed from the 12Mb yeast genome (sacCer3) in silico. Paired-end FASTQ sequences were randomly sampled assuming a uniform distribution from these in silico-modified genomes at varying levels of theoretical sequencing depth. The simulated FASTQ files were aligned using BWA-MEM to the appropriate reference genome (sacCer3). These BAM files and their respective species genomic interval information were fed to the DeletionID system. The correct deletion was reliably identified within the yeast genome using 3M unique paired-end tags 93.6% and 92.3% of the time for the simulated *reb1Δ* and *rap1Δ* knockouts respectively and improves with increased depth (**Figure 4A; Supplementary Table 5**). This represents the recommended minimum sequencing depth required for DeletionID to successfully identify the deletion interval in yeast whole-genome sequencing data. DeletionID leverages the efficient pysam library to parse BAM files, running to completion in under 1 minute on average for 3M reads of a yeast dataset (**Figure 4B**; **Supplementary Table 5**).

**Figure 4.**
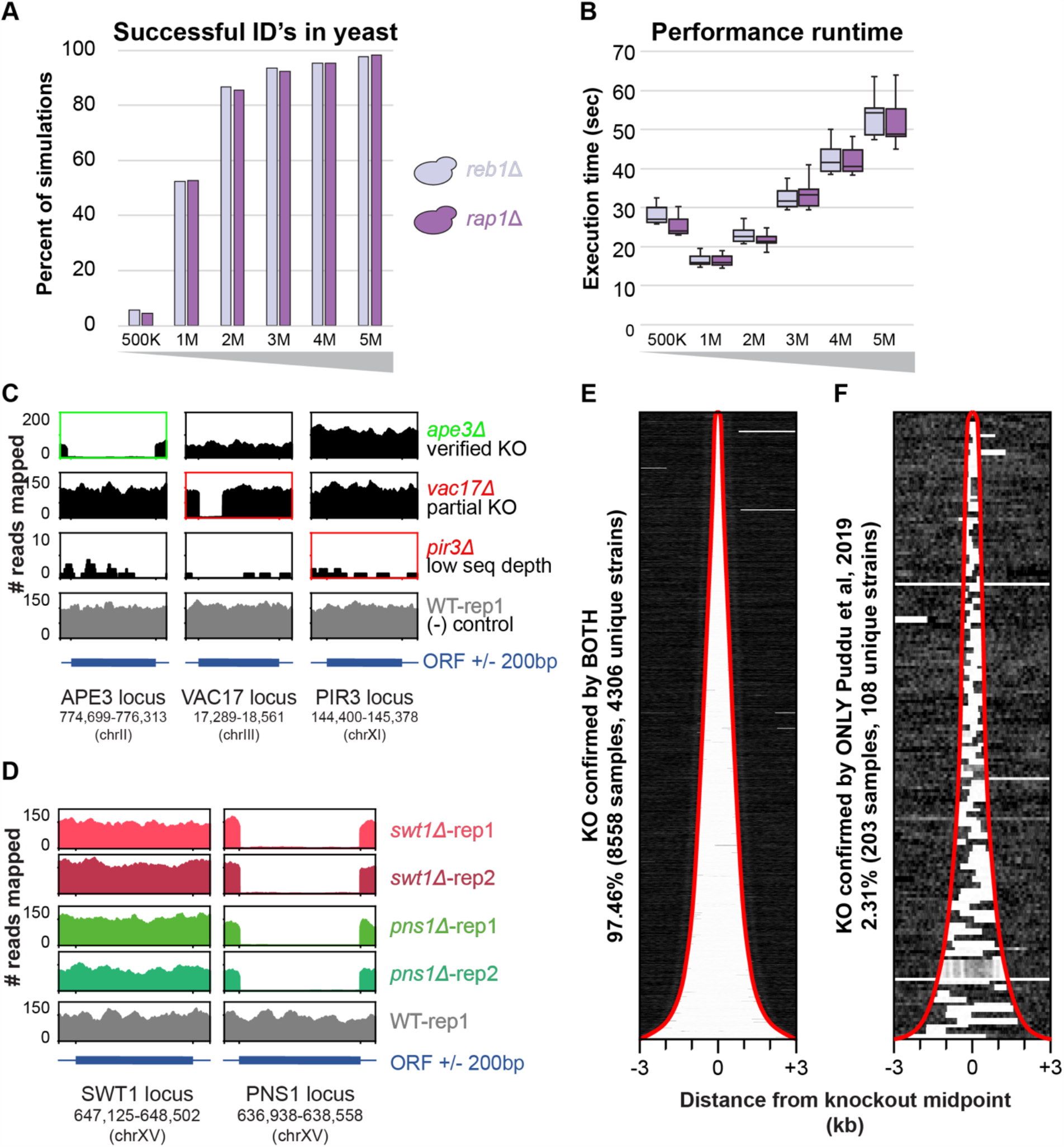
Evaluation of DeletionID for sensitivity, performance, and detection of knockouts in whole genome sequencing datasets. (**A**), Simulations for *reb1*Δ (red) and *rap1*Δ (yellow) genetic backgrounds show the number of simulated datasets that DeletionID successfully identified the deletion interval (y-axis) at each simulated dataset size (x-axis). (**B**), Box plot of runtime performance for simulations in seconds (y-axis) for each simulated dataset size (x-axis) of *reb1*Δ (red) and *rap1*Δ (yellow) genetic backgrounds. (**C**), Read coverage for three knockout samples and a control wild-type sample are shown across three loci (APE3, VAC17, and PIR3). The green text indicates a sample in which the expected knockout was confirmed by DeletionID and the red text indicates samples in which DeletionID did not identify the expected knockout. (**D**), Read coverage for four samples with *swt1*Δ and *pns1*Δ backgrounds (two replicates each) and a control wild-type sample are shown across two loci (SWT1 and PNS1). Despite the expected knockout interval around the SWT1 locus for the *swt1*Δ samples (first and second rows), no depletion is observed. The PNS1 locus, which was identified as a depleted interval in the DeletionID reports for the *swt1*Δ samples, shows a start-to-stop deletion of coverage that matches the coverage of the *pns1*Δ samples (third and fourth rows). (**E**,**F**), The normalized read coverage of the local interval around the center of the expected knockout (+/- 3 kilobases) for each sample (row) is shown in the two heatmaps with samples sorted by expected knockout interval length. The red traces mark the start and end intervals of the expected knockout regions. (**E**), Samples whose knockouts were confirmed by both Puddu et al, 2019 and DeletionID show mostly samples with knockouts concordant with their annotations. (**F**), Samples whose knockouts were confirmed by only Puddu et al, 2019, and not by DeletionID show the samples contain knockouts that are discordant with the expected knockout intervals.

### Verification of genomic deletions in YKOC knockout collection

The Yeast Knockout Collection (YKOC) is a collection of *S. cerevisiae* knockout strains that has been commonly used in many genetic studies and screens over the past two decades (57,58). A recent study performed whole genome sequencing (WGS) on ∼4,500 diploid strains from the collection that were reported to be homozygous for precise start-to-stop deletions of annotated ORFs in an effort to investigate the effects of these deletions on genome stability (51). We used this data to determine if DeletionID can recapitulate their findings by confirming the presence of the YKOC strain’s designated knockout genes. The data included 9,014 distinct genomic datasets that were sequenced from 4,530 deletion strains with one to five replicates each and four “wild type” control strains. Of the 8,761 samples Puddu et al. confirmed knockouts for, DeletionID identified the full ORF knockout in 8,558 samples (97.46%) and failed to identify the expected knockout in 203 samples (2.31%). 95 samples were not included because they were either controls, the data were unavailable, or their knockout interval was a not included in the DeletionID reference set as a deleted ORF, merged ORF, pseudogene, or transposable element gene (**Supplementary Table 6**). The genes that DeletionID failed to identify as a knockout were samples where the actual knockout interval was discordant with the annotations used or where the sample’s low sequencing coverage indicated a failed experiment (**Figure 4C**). For example, the *ape3Δ* dataset possessed a confirmed DeletionID knockout with reads strongly depleted across the entire *APE3* interval other gene intervals such as *VAC17* and *PIR3*. The *vac17Δ* dataset, for which DeletionID did not identify the expected knockout, showed read coverage depletion across only part of the ORF, pointing to discordant annotation of the knockout. The *pir3Δ* dataset in which DeletionID also failed to identify the expected knockout shows poor read coverage across all three regions, demonstrating the difficulty of confidently calling a depletion of reads across any interval in a dataset with low sequencing depth. All samples were constructed in a *leu2Δura3Δ* background and DeletionID identified both *ura3Δ* and *leu2Δ* in all but four low coverage samples (**Supplementary Table 6**).

DeletionID identified several potentially mislabelled samples like *swt1Δ* for which the DeletionID report indicated a *pns1Δ* background which was confirmed to match the *pns1Δ* datasets upon manual inspection of the alignments (**Figure 4D**). The Puddu et al. knockout-verified samples were successfully identified by DeletionID when the knockout annotation boundaries were concordant with the annotations used by DeletionID. In some samples where annotations were discordant, the interval checked by DeletionID fell within the actual knockout interval (**Figure 4E**). The datasets that DeletionID did not confirm the expected knockout for all appeared to have discordant intervals such that there was significant read coverage within the interval DeletionID checked. This coverage was sufficient for DeletionID to not mark it as an interval depleted of reads (**Figure 4F**). Small knockout interval lengths rendered these intervals more sensitive to being marked as not depleted because just a handful of reads could drive up the coverage of the interval much faster than if it was averaged across a longer interval.

While a confirmed deletion rate of >97% ostensibly seems ideal and is truly a testament to high-quality controls and laboratory practice, we note that the ∼2% failure rate for strains in the YKOC library still reflects over 200 genomic datasets that were not accurate as to what was published. Recent work has identified and corrected several of the YKOC strains we also identified in this study as wrong (58). However, this example clearly demonstrates the limitations of a purely biochemical quality control system and shows that DeletionID is capable of both confirming trust in existing samples and removing improper samples from downstream analysis.

### StrainID Module: Genomic variant identification

The StrainID module looks for the presence of strain-specific variants within HTS data to calculate a score for each variant profile (i.e., VCF) that measures how similar a dataset matches the variant profile (**Figure 1C**). The algorithm functions by searching aligned reads (i.e., BAM) to tally up those that match the alternate alleles and those that match the reference from the variant sites within each variant profile. From this a preliminary score for each profile is obtained. Then a background model is generated to account for the total genetic variation and error rates in the sample by approximating how much the sample’s reads deviate from the reference genome. This is calculated by randomly sampling reads from the dataset and performing another tally where deviations from the reference are counted and turned into a ratio of deviations from the genome over total sites checked. This background score is used to scale each variant profile score to determine which variant profile best matches the variant signals found in each dataset. Real datasets include some noise due to errors or spontaneous mutations that arise from genetic drift. ENCODE alignment data show that despite this noise, a dominant signal of the expected alternate allele is present in the reads from one background, such as HeLa, and absent in the reads from a different strain background, such as K562 (**Figure 5A**).

**Figure 5.**
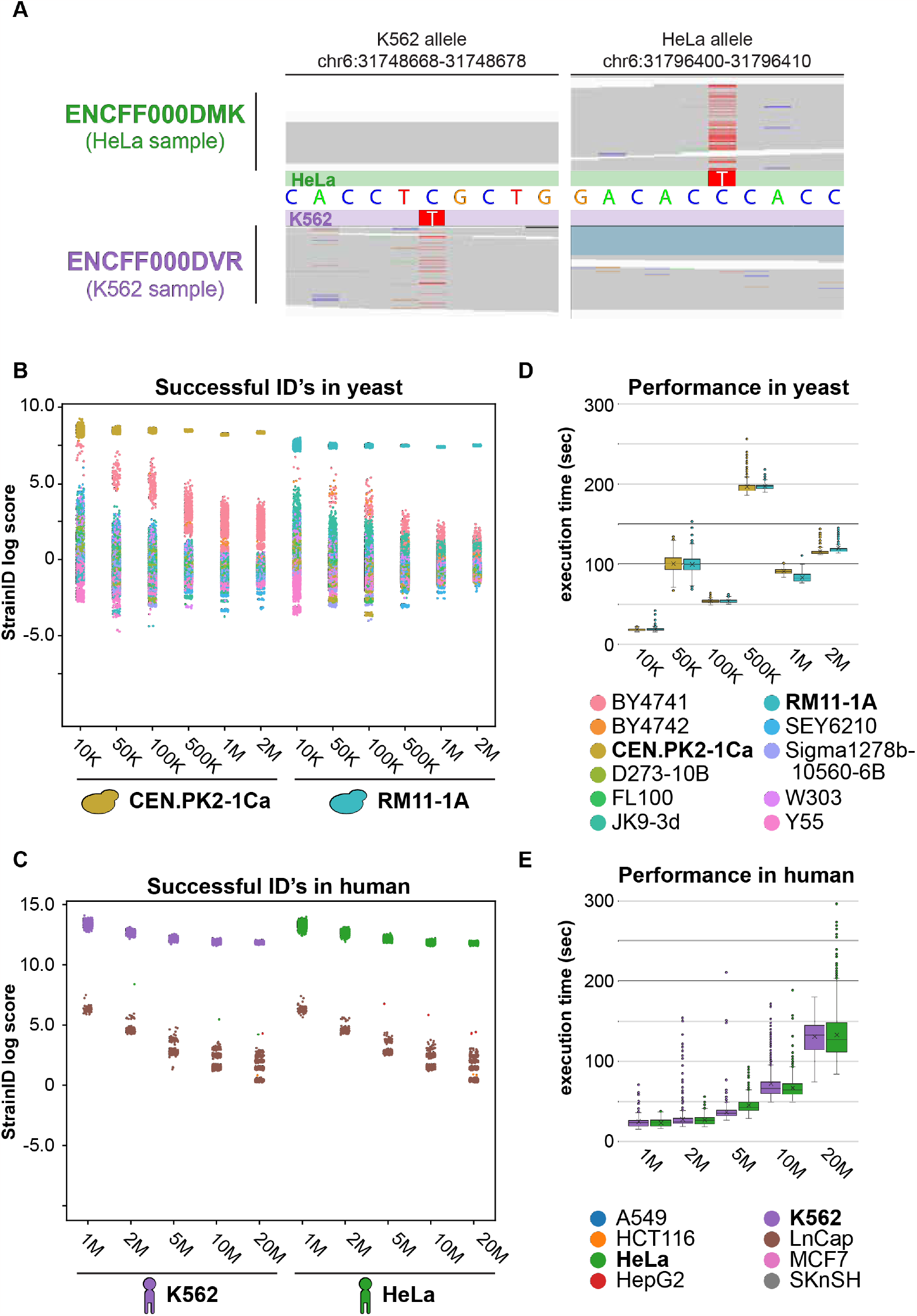
Evaluation of the StrainID module for sensitivity and performance. (**A**), Alignments for two ENCODE samples from HeLa (ENCFF000DMK) and K562 (ENCFF000DVR) genetic backgrounds are shown at two polymorphic loci. The first locus includes a known variant in the K562 background but not the HELA background and the second locus includes a known variant in the HELA background but not in the K562 background. (**B**), Simulated datasets were generated at 10K, 50K, 100K, 500K, 1M, and 2M depth from CENPK and RM11-1A backgrounds. StrainID scores (y-axis) for each variant profile/VCF (color) are grouped by sequencing depths and strain background (x-axis). (**C**), Human simulations were based on K562 and HELA backgrounds using a titration of 1M, 2M, 5M, 10M, and 20M simulated reads. (**D**),(**E**), Runtime performance of these simulations was measured in seconds and shown for both reference genomes (d, yeast; e, human).

The StrainID algorithm was developed by simulating HTS data from four in silico-modified genomes, each containing a different set of variant profiles over two organisms. The variant profiles used came from four VCF files capturing variants from CEN.PK2 and RM11-1A strains on the sacCer3 yeast genome build, and from K562 and HeLa cell lines on the hg19 human genome build. Randomly sampled datasets with sequencing depths of 10K, 50K, 100K, 500K, 1M, and 2M paired-end reads were constructed from each of the sacCer3 in silico-modified genomes and datasets with sequencing depths of 1M, 2M, 5M, 10M, and 20M were constructed from each of the hg19 in silico-modified genomes. These simulated datasets were all run through StrainID using the default variant database provided by GenoPipe for the appropriate genome assembly and matching reference FASTA file.

In our simulations, CEN.PK2 and RM11-1A simulations at 50K paired-end reads or more show good separation of the correct strain scores from the scores of the incorrect strains. This separation is determined by whether all correct strain scores for the simulations from a specific strain background and depth are greater than 2 standard deviations away from the average strain score. (**Figure 5B; Supplementary Table 7**). For the 50K yeast simulations, the threshold two standard deviations above the average are 7.89 and 6.37 for CEN.PK2 and RM11-1A simulations respectively. In the human simulations, the vast majority of scores (all from incorrect strains) were “NaN” or invalid numbers too be excluded from calculations. As a result, the standard-deviation based metric was adjusted so that the threshold is not driven primarily by correct strain values. Thus, for human simulations, the threshold was determined by two standard deviations above the average where both the standard deviation and the average were calculated excluding the correct strain scores. With this threshold, we establish that 1M paired-end reads is sufficient for reliable detection of the correct strain using StrainID (**Figure 5C; Supplementary Table 8**). For both yeast and human, the separation of the correct score from the other strain scores increased with increased read depths, showing that more reads can improve the accuracy of the results and the increasingly tighter distributions of the correct strain scores indicates an improvement in precision as well. In addition to having a high specificity for identifying the correct strain, StrainID’s runtime performance is short because it is not attempting to identify variants in a de novo fashion, but rather leverages the efficient BAM file parsing library, pysam, to search for known variants and calculate a score reflecting how well the variant profile of the data matches the known strain profile (47). The runtime at recommended 1M dataset sizes for yeast and human using the GenoPipe provided default variant databases is approximately 90 seconds and 30 seconds for yeast and human respectively (**Figure 5D-E; Supplementary Table 7**,**8**). All simulations showed 100% of the datasets assigned the correct strain with the best score, however real data can be more complicated due to the introduction of noise and non-uniform coverage, so StrainID’s performance was further evaluated in real datasets.

### Genomic variant identification within real yeast datasets

To demonstrate StrainID’s utility in real yeast datasets, we ran StrainID on published yeast ChIP-seq datasets from CEN.PK and BY4742 backgrounds (52,53).The default variant database for the sacCer3 genome build includes VCF files that are subsets of the complete set of variants. The variants kept are unique to their respective strains and not found in any other VCF files to ensure the specificity of the variants checked by StrainID. For example, from the complete set of BY4742 variants, those variants shared with BY4741, or any other strain background are removed (**Figure 6A**). StrainID confidently identified the background of the CEN.PK samples with significant separation of the correct CEN.PK score from the scores of the other strains. In eight of the ten BY4742 samples, StrainID identified BY4742 as the strain background with the best match to the data (**Figure 6B; Supplementary Table 9**). For the two samples that did not identify BY4742, the highly genetically-similar BY4741 strain was identified as the best strain match. Furthermore, in all samples both BY4742 and BY4741 scores showed high separation from those of other strains, adding to the confidence in calling the background of the BY4742 samples as from at least one of the BY strain backgrounds.

**Figure 6.**
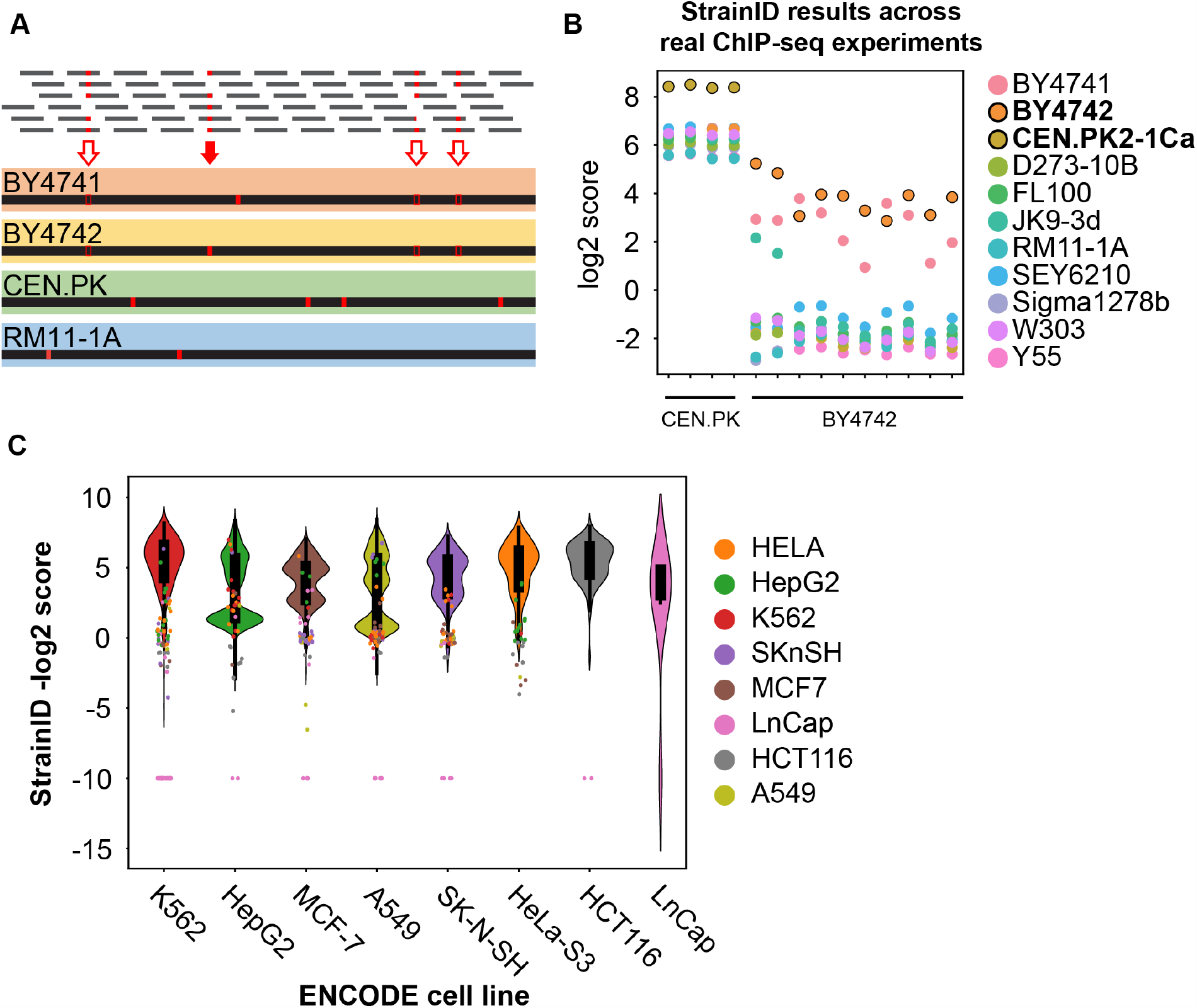
The StrainID module consistently and reliably identifies the correct strain/cell line in real datasets. (**A**), The yeast variant database for StrainID was built from full sets of variants for each strain with the red arrows marking the variants for BY4742 (top) and then filtered to only include unique variants with the red arrows indicating the variants kept in the BY4742 strain (bottom). (**B**), StrainID scores (y-axis) for each variant profile checked (colour) are shown for each dataset (x-axis) to show the separation of the correct strain score from the other strains. (**C**), The StrainID scores for several thousands of ENCODE samples are displayed such that successfully identified backgrounds are summarized by the violin plots while the scattered points indicate samples that were misidentified, as they appear to be from a different strain background indicated by the coloured dots. The coloured dots are more closely linked to their same-colour violins.

## DISCUSSION

Confidence in the results of an experiment is dependent on the veracity of the sample metadata. However, error can be introduced at nearly every stage of the experimental process. Previous attempts to verify the cellular identity of genomic datasets have been limited to a few specific use-cases. Additionally, the current gold-standard for validating cellular identity (STR profiling) does not necessarily preclude downstream sample mix-up. It also does not validate the identity of CRISPR modified cell lines that may be genotypically distinct relative to its source cell line while still possessing an identical STR profile. To provide a mechanism by which the identity of cellular material can be confirmed post-sequencing, we developed GenoPipe. The three core modules of GenoPipe: EpitopeID, DeletionID, and StrainID were developed to identify major genotypical determinants of cellular identity. We demonstrated that GenoPipe can detect genotype perturbations at realistic and practical sequencing depths as defined by ENCODE (14) and that the execution speed of each module was reasonable (seconds to minutes) using standard computational hardware. We successfully validated GenoPipe across a variety of organisms using a broad range of epigenomic assays (e.g., ChIP-seq, WGS, ATAC-seq).

As epitope-tagging of protein becomes more common in genomics research, particularly due to the difficulties in acquiring native antibodies, opportunities for sample mislabelling also increases. Large consortium projects such as ENCODE have adopted tagging techniques to study eukaryotic genomes, generating thousands of unique datasets in genetically modified cell lines. We found that while the vast majority of datasets generated by ENCODE in these lines were labelled correctly, several datasets were called by GenoPipe as possessing inconsistent sample metadata and should be investigated more deeply. Some of these inconsistencies may be attributed to the read-hopping phenomenon found in data generated from sequencing platforms using pattern flow-cell technology while others appear to be a more clear-cut case of sample mislabelling (59). Increased adoption of paired-end sequencing will also result in the ability to localize epitope-tags within strains as well demonstrating an additional value-add for paired-end HTS technology. EpitopeID’s identification of common HIV integration sites also provide an additional application of the module that can be further developed for sensitive detection of structural rearrangements like common translocations.

In addition to identifying the presence and location of synthetic protein epitopes, we also used GenoPipe to examine the accuracy of the Yeast KnockOut Collection (YKOC), a collection of thousands of unique yeast strains possessing specific gene knockouts that has been maintained for decades. Over the years, multiple mistakes in the collection have been identified and corrected from the original YKOC resulting in updated collections (50). We tested the original collection sourced by Puddu et al. from EUROSCARF in 2001 and GenoPipe identified several strains with incomplete or truncated gene knockouts that did not match the YKOC annotations as well as potentially mislabelled strains that warrant further biochemical investigation. GenoPipe also identified samples with partial coverage within the expected genomic interval at approximately half the global coverage depth. This may indicate a potential erroneous cross with another strain resulting in a heterozygous knockout or simple contamination of the stock by another strain and demonstrates GenoPipe’s potential to identify heterozygous imbalances in diploid organisms.

The vast body of data generated by ENCODE is based on common cell lines with unique and known variant profiles. Like EpitopeID and DeletionID, StrainID may be of particular interest to consortiums generating large amounts of HTS data on various cell lines especially in light of the well-documented issues of cell line contamination in research. Using StrainID on a selection of ENCODE-derived data we identified several potentially contaminated samples that may need to be replicated to check for contamination. The samples with weak StrainID scores from assays such as single end small-RNA and CAGE would likely perform better with a different set of variants that include those that fall in intergenic and promoter regions to increase the coverage across the SNP sites.

The flexible construction of GenoPipe allows for the expansion of GenoPipe to quality control any organism with a reference genome and any genomic assay that generates background at a sufficient depth. Genome complexities such as high ploidy in plants or high paralogous genes in Zebrafish present unique challenges that can be addressed in future versions of GenoPipe. Incorporating additional known genotype information such as copy number variation will also likely improve the accuracy of GenoPipe in a wide variety of samples. However, in its current form, GenoPipe can easily be integrated as a single step quality control verification of the sample’s genetic background for laboratories generating sequencing data and provides an additional level of quality control and data confidence. We envision GenoPipe to become a valuable and necessary tool in the toolkit for quality control of HTS data.

## AVAILABILITY

GenoPipe is an open-source tool available on Github (https://github.com/CEGRcode/GenoPipe). The scripts used for benchmarking and testing in this manuscript may also be found in this repository.

## ACCESSION NUMBERS

Nonapplicable

## ACKNOWLEDGEMENT

The authors thank the members of Pugh lab whose work made this pipeline required. We thank the members of the Cornell EpiGenomics Core (EGC, RRID:SCR_021287) for their feedback and contributions over the course of this project.

